# Probing the Acyl Carrier Protein-Enzyme Interactions within Terminal Alkyne Biosynthetic Machinery

**DOI:** 10.1101/291195

**Authors:** Michael Su, Xuejun Zhu, Wenjun Zhang

**Affiliations:** Department of Chemical and Biomolecular Engineering, University of California, Berkeley, 2151 Berkeley Way, Berkeley, CA 94720; Chan Zuckerberg Biohub, San Francisco, CA 94158

## Abstract

The alkyne functionality has attracted much interest due to its diverse chemical and biological applications. We recently elucidated an acyl carrier protein (ACP)-dependent alkyne biosynthetic pathway, however, little is known about ACP interactions with the alkyne biosynthetic enzymes, an acyl-ACP ligase (JamA) and a membrane-bound bi-functional desaturase/acetylenase (JamB). Here, we showed that JamB has a more stringent interaction with ACP than JamA. In addition, site directed mutagenesis of a non-cognate ACP significantly improved its compatibility with JamB, suggesting a possible electrostatic interaction at the ACP-JamB interface. Finally, error-prone PCR and screening of a second non-cognate ACP identified hot spots on the ACP that are important for interacting with JamB and yielded mutants which were better recognized by JamB. Our data thus not only provide insights into the ACP interactions in alkyne biosynthesis, but it also potentially aids in future combinatorial biosynthesis of alkyne-tagged metabolites for chemical and biological applications.

**Topical Heading:** Biomolecular Engineering, Bioengineering, Biochemicals, Biofuels, and Food

## Introduction

The alkyne (C≡C) is a readily derivatized functionality valued for its diverse applications in nearly all areas of modern chemistry and biology. It is widely employed in the alkyne-azide Huisgen cycloaddition reaction (known as a click reaction), which has allowed modular assembly of complex chemical structures for both drug development and material synthesis.^1^ In addition to its chemical applications, the alkyne is also an important bio-orthogonal tag in chemical biology. The alkyne can be metabolically or chemically installed into medicinally and biologically important molecules, such as glycans, proteins, nucleic acids, lipids, and natural products; and the mode of actions and cellular dynamics of these alkyne-tagged molecules can be further investigated through click chemistry to introduce an analytical handle,^2,3^ or directly using stimulated Raman scattering microscopy.^4^ Considering the importance of alkynes in both chemistry and biology, a synthetic biology route for alkyne synthesis and installation is highly desired.

In contrast to the broad applications of alkynes, few biosynthetic enzymes have been characterized for alkyne synthesis. We recently elucidated an acyl carrier protein (ACP)-dependent terminal alkyne biosynthetic pathway in which a fatty acyl-ACP ligase activates a fatty acid using ATP and further attaches it to a cognate ACP, followed by the sequential terminal dehydrogenation of the fatty acyl moiety catalyzed by a membrane-bound bi-functional desaturase/acetylenase. In particular, two homologous pathways, JamABC from *Moorea producens* JHB and TtuABC from *Teredinibacter turnerae* T7901, have been identified which activate hexanoic acid and decanoic acid, respectively. We further showed that omitting the ACP, either JamC or TtuC in the respective pathway, completely abolished or significantly impaired alkyne synthesis,^5,6^ demonstrating the necessity of ACP in terminal alkyne biosynthesis.

Not only critical to alkyne biosynthesis, carrier proteins (CPs) in general are central hubs in the biosynthesis of fatty acids and medicinally important polyketide (PK) and non-ribosomal peptide (NRP) natural products^7^ (**Figure 1**). Selective and programmed CP recognitions by multiple biosynthetic enzymes are essential for the precise assembly of complex natural product scaffolds, and combinatorial biosynthesis using simple mix-and-match strategy often yields significantly impaired assembly lines due to CP incompatibility. Thus, there is a crucial demand to understand CP interactions with their catalytic partners. While some information about transient CP-enzyme interactions is emerging via X-ray crystallography,^8,9^ NMR,^10^ electron cryo-microscopy (cryo-EM),^11^ and crosslinking technology,^8,12^ how ACP selectively interacts with alkyne biosynthetic enzymes, particularly the fatty acyl-ACP ligase and the bi-functional desaturase/acetylenase, is not well understood. Preliminary studies of the substrate specificity of JamB, the acetylenase/desaturase, implied strong protein-protein interactions between the acetylenase/desaturase and the dedicated ACP,^5^ warranting the further investigation of the central role of ACP in alkyne synthesis. The flexibility of ACP in the alkyne biosynthetic machinery is particularly important for the application of these enzymes in engineered biosynthetic pathways to generate and incorporate this functionality into various PK/NRP scaffolds. Here we try to extend our insights into ACP recognitions in terminal alkyne biosynthesis through protein engineering using JamABC as a model system.

**Figure 1.**
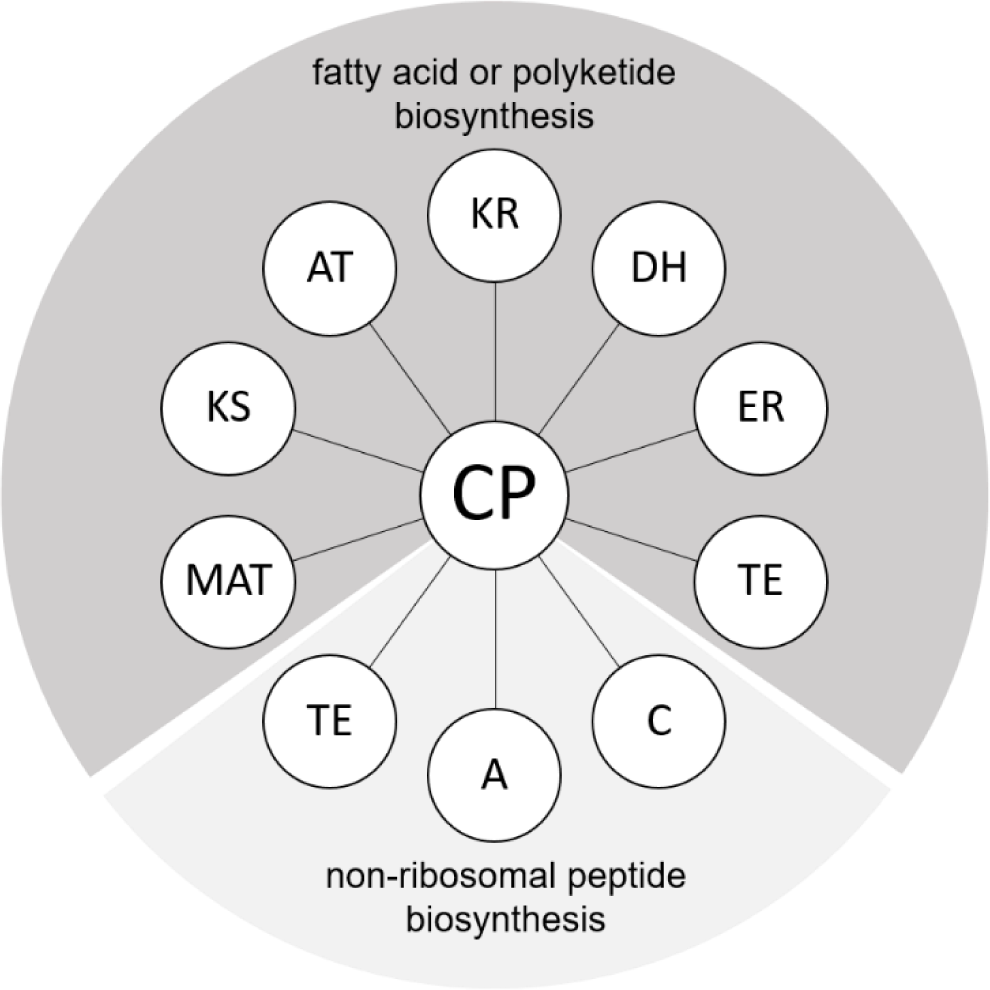
Carrier proteins interact with different catalytic partners involved in fatty acid, polyketide, and non-ribosomal peptide biosynthesis. MAT: malonyl-CoA:ACP transferase; KS: ketosynthase; AT: acyl transferase; KR: ketoreductase; DH: dehydratase; ER: enoyl reductase; TE: thioesterase; C: condensation domain; A: adenylation domain.

## Materials and Methods

### Bacterial Strains and Plasmid Constructions

The full names of the ACP genes used in this study as well as the DNA templates used to PCR amplify them are listed in **Table S1**. Plasmid constructions were performed using standard protocols. Site-directed mutagenesis was performed using the standard QuikChange strategy using relevant templates. Plasmids were purified from *E. coli* XL1-Blue with Zyppy™ Plasmid Miniprep Kit and confirmed by DNA sequencing (UC Berkeley DNA Sequencing Facility). *E. coli* BAP1^13^ and *E. coli* BL21 Gold (DE3) were used for protein expression. All primers used in this study were ordered from Integrated DNA Technologies. Complete lists of plasmids and primers used in this study are provided in **Tables S2 and S3**.

### Protein Expression and Purification

All ACPs purified in this work contained C-terminus hexahistidine tags and their expression plasmids were transformed into *E. coli* BAP1. The cells were grown at 37 °C in 750 mL of LB medium with appropriate concentrations of antibiotics to an OD_600_ of 0.4-0.6, followed by the induction with 0.12 mmol/L isopropyl-β-d-thiogalactopyranoside (IPTG) for 16 h at 16 °C. The cells were harvested by centrifugation (4424 g, 15 min, 4 °C), resuspended in 30 mL of lysis buffer (50 mmol/L HEPES, pH 8.0, 300 mmol/L NaCl, 10 mmol/L imidazole), and lysed by homogenization on ice. Cell debris was removed by centrifugation (15000 g, 1 h, 4 °C). Ni-NTA agarose resin was added to the supernatant (1.5 mL/L of culture), and the solution was nutated at 4 °C for 1 h. The protein–resin mixture was loaded onto a gravity flow column, and proteins were eluted with increasing concentrations of imidazole in buffer A (50 mmol/L HEPES, pH 8.0, 300 mmol/L NaCl). Purified proteins were concentrated and buffer exchanged into HEPES buffer (50 mmol/L HEPES, pH 8.0, 100 mmol/L NaCl) with Amicon Ultra centrifugal filters. The final proteins were flash-frozen in liquid nitrogen and stored at −80 °C. Protein concentrations were determined by NanoDrop. The fatty-acyl ACP ligase JamA was expressed and purified as previously described.^5^

### *In vitro* ACP Loading Assays

Assays were performed in 50 μL of 50 mmol/L HEPES (pH 8.0) containing 2 mmol/L MgCl_2_, 2 mmol/L ATP, 1 mmol/L TCEP, 5 mmol/L fatty acids, and 50–100 μmol/L holo-ACP. The reactions were initiated by the addition of 10 μmol/L JamA and incubated at room temperature. At regular time intervals (5, 10, 20 min), 10 μmol/L aliquots were quenched with 10 μL of 10% (v/v) formic acid. The mixture was diluted two-fold, and centrifuged at 4 °C to remove any precipitate. 10 μL of the resulting protein sample was analyzed by liquid chromatography-high resolution mass spectroscopy (LC-HRMS) with an Aeris 3.6 μm widepore XB-C18 column (250 × 2.1 mm). A linear gradient of 15–98% CH_3_CN (v/v) over 16 min and 98% CH_3_CN for a further 20 min in water supplemented with 0.1% (v/v) formic acid at a flow rate of 0.15 mL/min was used for analysis. MassHunter Qualitative Analysis software was used for data analysis, and the intact protein masses were obtained using a maximum entropy deconvolution algorithm from Agilent MassHunter BioConfirm Software. The parameters used in this algorithm: a mass range of 9000-17000 Daltons, a mass step of 1.0 Dalton, a signal-to-noise peak threshold of 30, a proton-based adduct, a 90% average mass peak height, an automatic isotope width, a minimum consecutive charge state of 5, and a minimum protein fit score of 8.

### Biosynthesis of Terminal Alkyne-tagged Polyketides in *E. coli*

*E. coli* BAP1 strains were transformed with plasmids containing *jamA, jamB, hspks1*, and the gene encoding the appropriate ACPs. A BAP1 strain co-expressing *jamABC* and *hspks1* was developed in each *in vivo* experiment as a positive control. **Table S4** provides a complete list of developed *E. coli* BAP1 strains used in this study. The *E. coli* cells were grown in 200 mL of LB medium with 100 mg/L carbenicillin and 100 mg/L spectinomycin at 37 °C to an OD_600_ of 0.4–0.45. Subsequently, these cells were harvested and concentrated 5-fold into 40 mL of fresh F1 medium (1L contains 3 g KH_2_PO_4_, 6.62 g K_2_HPO_4_, 4 g (NH_4_)_2_SO_4_, 150.5 mg MgSO_4_, 5 g glucose, 1.25 mL trace metal solution, 100 μmol/L Fe(NH_4_)_2_(SO_4_)_2_, and 10 mmol/L 100x vitamin solution) supplemented with 100 mg/L carbenicillin, 100 mg/L spectinomycin, 0.6 mmol/L IPTG, and 1 mmol/L 5-hexenoic acid. After 48 hours of growth and expression at 20 °C, compounds were extracted from cell-free supernatant with ethyl acetate. The solvent was removed by rotary evaporation, and the residue was redissolved in methanol and analyzed by LC-HRMS with an Agilent Eclipse Plus C18 column (4.6 × 100 mm). A linear gradient of 20–50% CH_3_CN (v/v) over 15 min, 95% CH_3_CN for a further 3 min, and 20% CH_3_CN for a final 5 min in H_2_O supplemented with 0.1% (v/v) formic acid at a flow rate of 0.5 mL/min was used for LC-HRMS analysis. LC-HRMS analysis was performed on an Agilent Technologies 6510 Accurate Mass QTOF LC-MS.

### Error-Prone PCR and Construction of *peACP* Mutant Library

The error-Prone PCR procedure was modified from established protocols.^14^ The reaction consisted of 0.35 mmol/L dATP, 0.4 mmol/L dCTP, 0.2 mmol/L dGTP, 1.35 mmol/L dTTP, 1.75 mmol/L MnCl_2_, and 1 U Taq polymerase. The reaction mixture was submitted to 25 cycles of PCR: 95 °C for 1 min, 50 °C for 1 min, and 68 °C for 1 min. The resulting PCR products were digested with NcoI/BamHI and inserted via cohesive end ligation into pXZ34, a pETDuet-1 plasmid containing *jamB*, that was pre-digested with NcoI/BamHI. The cohesive end ligation was executed by following the NEB Quick Ligation Kit’s protocol. The ligation product was purified using spin column centrifugation, co-transformed with pXZ27, a pCDFDuet-1 plasmid containing *jamA* and *hspks1*, to an electrocompetent BAP1 strain, and plated on LB agar containing 100 mg/L carbenicillin and 100 mg/L spectinomycin.

### Plate-Reader Fluorescence Measurements

The measurements were performed following a modified protocol established previously.^15^ Reactions were run in a 96-well black plate, and measurements were performed using a Tecan Safire Microplate Reader. Each well had a 200 μL sample containing 2.25 μmol/L of azido probe di-pegOF, 5 mmol/L ascorbic acid, 500 μmol/L CuSO_4_, 100 μmol/L BTTAA, and 190 μmol/L of cultures. The reactions were performed in dark for 10 min at room temperature.

### Bioinformatic Analysis of Proteins

Amino acid sequence alignment of the ACPs was performed using the online program Clustal Omega.^16^ Protein structures of JamB and the ACPs were modeled using the online program HHPred^17^ on the basis of a membrane-bound desaturase SCD1 (PDB: 4ZYO/4YMK)^18,19^ and CurA-ACP (PDB: 2LIU),^10^ respectively. ACP-JamB models were created by docking the ACP to the electropositive surface of JamB near the substrate entrance using ClusPro.^20,21^ The visualizations of each structure and model were created using the UCSF Chimera package. Chimera is developed by the Resource for Biocomputing, Visualization, and Informatics at the University of California, San Francisco.^22^

## Results and Discussion

### Studying the ACP Recognition by Acyl-ACP Ligase JamA

We first selected 15 ACPs from different natural product biosynthetic pathways to study the ACP recognition by JamA, the fatty acyl-ACP ligase. The ACPs we tested included JamC, seven JamC homologs^6^ (TtuC, BpACP, PeACP, CyACP, PfACP, IlACP, SpACP) found in putative alkyne biosynthetic machinery, and seven ACPs involved in the biosynthesis of erythromycin (DEBS1-ACP_L_),^23^ daptomycin (DptF),^24^ mycosubtilin (MycA-ACP_L_),^25^ A54145 (LptF),^26^ CDA (Sco3249),^27^ isonitrile lipopeptide (ScoB),^28^ and antimycin (AntD-ACP),^29, 30^ respectively. The detailed information of these ACPs is provided in **Table S1**. We first cloned each individual ACP-encoding gene into an expression vector that encoded a *C*-terminal His_6_-tag. The ACPs were then overexpressed in the *E. coli* BAP1 strain to ensure their posttranslational modifications to holo forms^13^ (**Figure S1**). The ratios of the holo and apo forms that coexist for each purified ACP used in the study are listed in **Table S5**. We next purified JamA from *E. coli* and tested its abilities to activate and load 5-hexenoic acid onto different ACPs using *in vitro* ACP loading assays, followed by liquid chromatography–high-resolution mass spectrometry (LC-HRMS) analysis (**Figure S2**). Approximately 25% of JamC was converted to 5-hexenoyl-JamC after 5 minutes of the *in vitro* reaction with JamA, and this conversion was set to be a relative JamA activity of 100%. The results suggested that JamA has relaxed substrate specificities with ACPs from the putative alkyne biosynthetic machinery: ScoB and all of the seven tested JamC homologs could be recognized by JamA to a varying degree (**Figure 2**). However, most of the tested ACPs from PK/NRP biosynthetic machinery could not be efficiently recognized by JamA. In particular, four ACPs exhibited a high degree of compatibility with JamA (TtuC, CyACP, BpACP, and PeACP with JamA activities of ~37%, ~41%, ~207%, and ~217% respectively), and they were therefore chosen for the study of protein-protein interactions with the subsequent enzyme JamB, the membrane-bound bi-functional desaturase/acetylenase.

**Figure 2.**
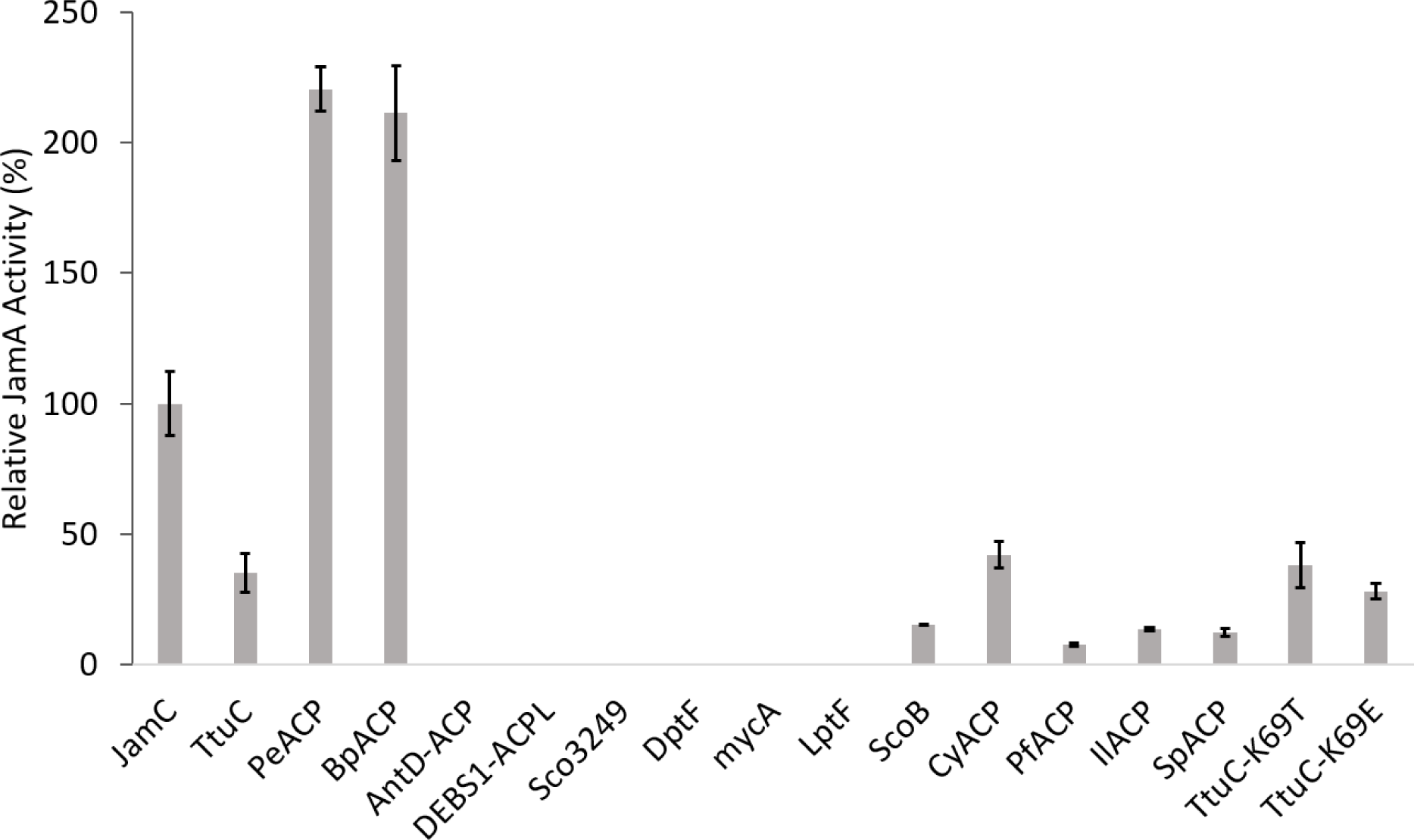
Relative JamA *in vitro* activities towards different ACPs. Approximately 25% of JamC was converted to 5-hexenoyl-JamC after 5 minutes of the *in vitro* reaction with JamA, and this conversion was set to be a relative JamA activity of 100%. Error bars represent standard deviations from at least three experiments.

### Studying the ACP Recognition by Acetylenase JamB

Due to the difficulties in working with membrane-bound proteins *in vitro*, we probed the ACP interactions with JamB *in vivo.* We previously reported that a promiscuous type III polyketide synthase (PKS), HsPKS1, could incorporate a carrier protein-bound fatty acyl moiety into the polyketide backbone, and consequently generate a terminal alkyne-tagged polyketide 1 in an engineered *E. coli* BAP1 strain that co-expressed *jamABC* and *hspks1* (named as strain XZ1) (**Figure. 3 and S3**).^5^ Since HsPKS1 shows no preference towards different carrier protein-bound acyl moieties,^6,15,31^ this *E.coli-hspks1* platform could be a feasible reporting system for the study of JamB-ACP interactions. We introduced ACP-encoding gene into the engineered *E. coli* BAP1 strain that co-expressed *hspks1, jamA*, and *jamB.* The resulting *E. coli* strains were grown in shake flasks at 20 °C for ~2 days with the supplementation of IPTG and 5-hexenoic acid. Upon the coexpression with *jamC, ttuC*, or *cyACP*, terminal alkyne-tagged polyketide **1** could be detected in the culture extracts by LC–HRMS analysis. In addition, the fed alkene precursor also led to a major byproduct 2 with a terminal alkenoyl modification (**Figure 3**). The compounds **1** and **2** were produced in an approximately 1:5 ratio by the strain XZ1,^5, 15^ and this efficiency was set to be a relative JamB activity of 100%. JamB activities towards TtuC and CyACP were ~39% and ~38%, respectively (**Figure 4**). No reliable production of **1** was observed from the culture extracts of the strains that co-expressed *bpACP or peACP*, suggesting weak/no recognition of JamB with these two ACPs (**Figure 4**).

**Figure 3.**
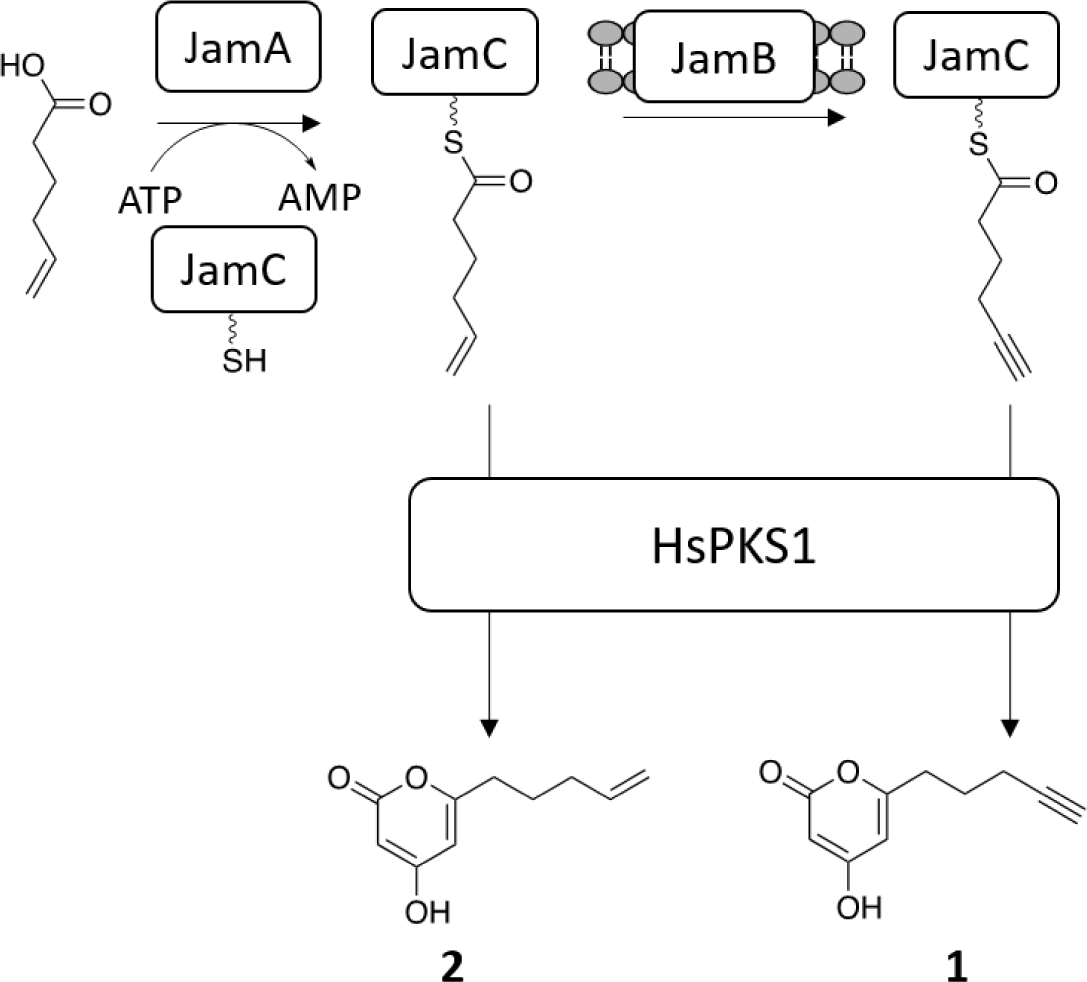
*E. coli-hspks1* platform for terminal alkyne biosynthesis. The balls/sticks on JamB indicate that it is a transmembrane protein. The promiscuous type III polyketide synthase, HsPKS1, incorporates a carrier protein-bound fatty acyl moiety into the polyketide backbone, and consequently generates a terminal alkyne-tagged polyketide **1** in an engineered *E. coli* BAP1 strain that co-expressed *jamABC* and *hspks1.*

**Figure 4.**
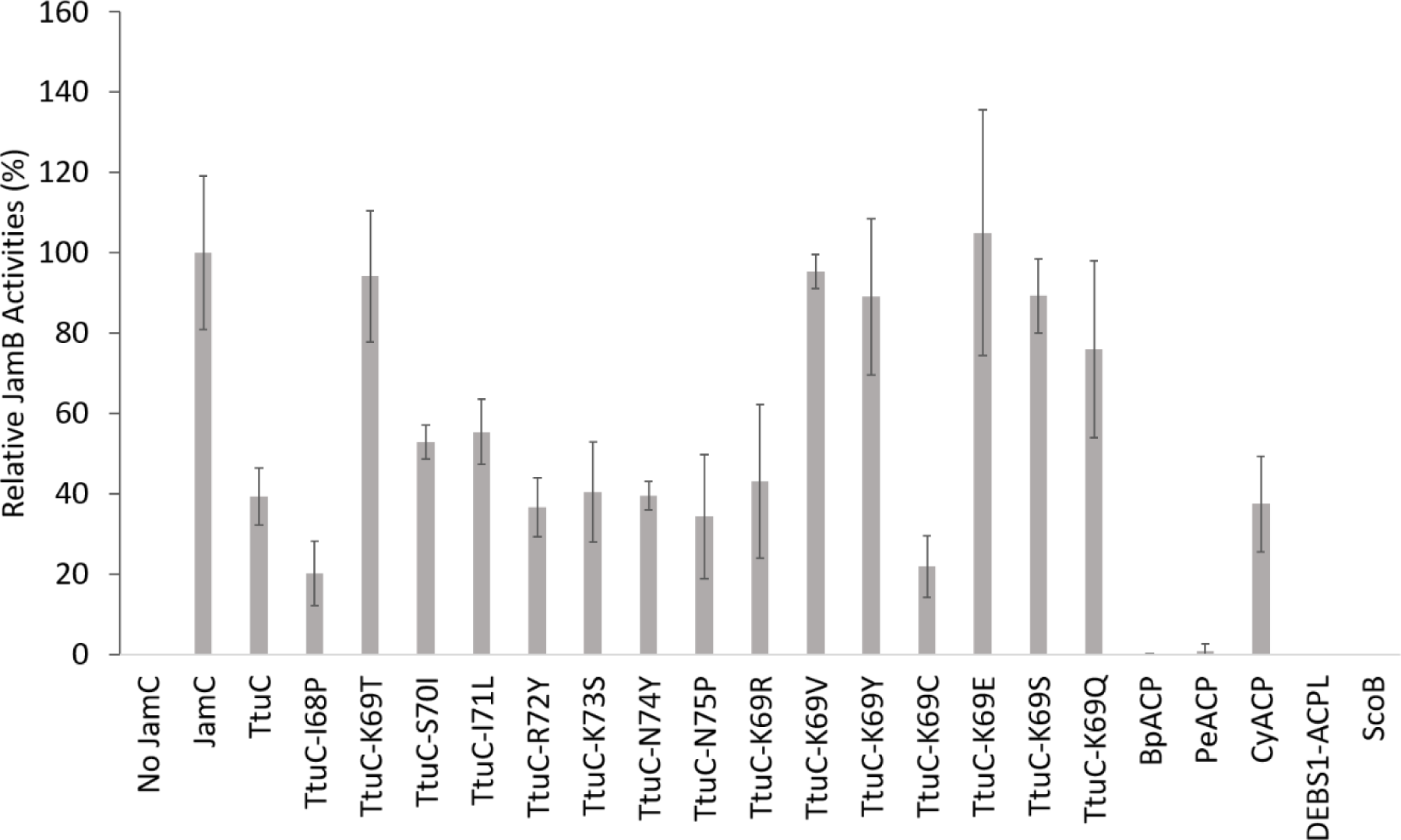
Relative JamB *in vivo* activities towards ACPs. The compounds **1** and **2** were produced in an approximately 1:5 ratio by the strain XZ1, and this efficiency was set to be a relative JamB activity of 100%. Error bars represent standard deviations from at least three experiments.

### Rational Engineering to Improve ACP Recognition by Acetylenase JamB

Due to the weak or no recognition of JamB with non-cognate ACPs, we next tried to test if protein engineering of ACPs could improve their interactions with JamB. The crystal structure of a soluble stearoyl-ACP desaturase in complex with an ACP (PDB: 2XZ0 and 2XZ1) revealed predominant electrostatic interactions between the two proteins: ACP contributed negatively charged residues while the soluble stearoyl-ACP desaturase displayed positively charged residues at the interaction surface.^32^ Despite the membrane-bound nature of JamB, we hypothesized that electrostatic interactions might contribute to JamB-JamC interactions. Our structural modeling of JamB and JamC and protein-protein docking analysis suggested electrostatic interactions at the JamB-JamC interface.^15^ To test this hypothesis, we set out to rationally engineer TtuC, BpACP, and PeACP by changing their electrostatic surfaces in an attempt to improve their interactions with JamB. Structural and sequence comparisons between TtuC and JamC revealed a positively-charged eight-amino acid region on TtuC helix 3 while the corresponding region of JamC lacks these positively-charged residues (**Figure. 5 and S4**). Individual amino acids in this region of TtuC were then mutated to the corresponding JamC amino acids using site-directed mutagenesis. The resulting *ttuC* mutants were then introduced into the *E. coli-hspks1* system to investigate their effects on compound **1** production. The introduction of K69T into *ttuC* led to an engineered *E. coli* strain (named as strain XZ2_2) with the compound **1** production efficiency comparable to XZ1 (expressing *jamC*); while the other mutations had no obvious effect (**Figure 4**). Since the K69T mutation had no effect on the TtuC expression or recognition by JamA based on SDS-PAGE and biochemical analysis (**Figure. 2 and S1**), we conclude that the improved alkyne production efficiency by XZ2_2 was due to the improved interaction between TtuC-K69T and JamB.

**Figure 5.**
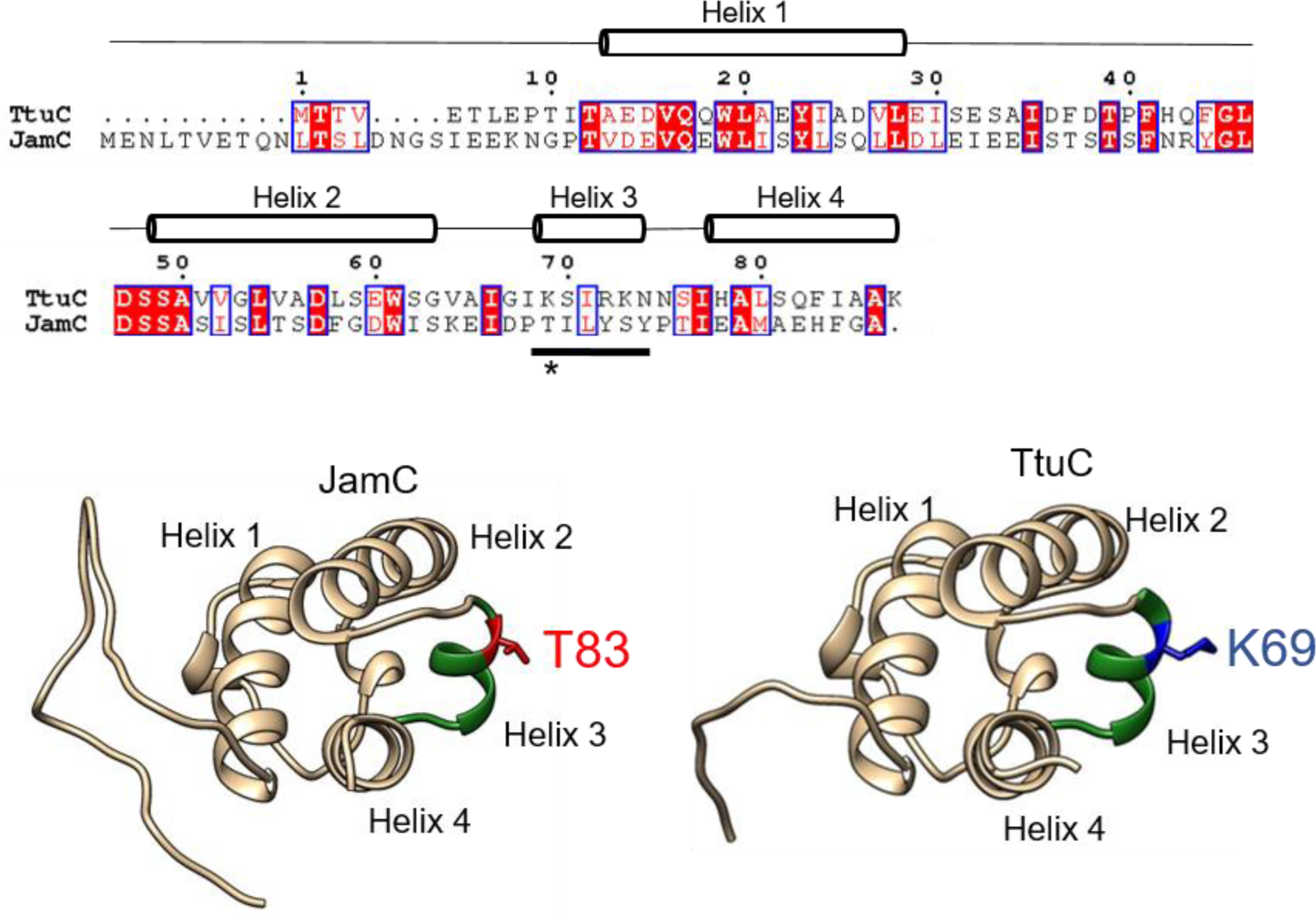
Comparison of JamC and TtuC. Top: The amino acid sequence alignment of TtuC and JamC. The helical regions predicted from the modelled structures are indicated. The residues on TtuC that were mutated in this study are underlined. TtuC’s K69 residue is indicated with an asterisk. Bottom: Modelled ribbon diagrams of JamC and TtuC. JamC-T83 and TtuC-K69 are shown as sticks.

To better understand the effect of amino acids with different properties at K69 of TtuC, we generated additional single mutants of TtuC, including K69R, K69E, K69Q, K69S, K69C, K69V, and K69Y, through site-directed mutagenesis. Most of these mutants showed improved alkyne production efficiency and as expected, TtuC-K69R had no significant improvement towards alkyne production compared to the wild-type TtuC (**Figure 4**). Out of all of the single mutants, the TtuC-K69E mutant had the highest alkyne production efficiency, with SDS-PAGE and biochemical analysis of this mutant indicating that this mutation also had no obvious effect on the TtuC expression or recognition by JamA (**Figure. 2 and S1**). These findings supported the sensitivity of JamB activity towards the electrostatic surface of TtuC and the critical role of the residue K69 on the Helix 3 of TtuC for interacting with JamB. Using a similar engineering strategy, however, we failed to improve the recognition of JamB with BpACP and PeACP; it is likely that engineering the electrostatic interactions were not sufficient for proper interactions between JamB and these ACPs.

### Error-prone PCR to Improve ACP Recognition by Acetylenase JamB

We then turned to the error-prone PCR and screening approach to evolve a heterologous ACP for improved recognition by JamB. This approach has previously been employed to rapidly improve the activity of chimeric assembly-line enzymes, including the recognition of non-cognate ACPs.^33,34^ PeACP was chosen as the target for error-prone PCR since it could be readily loaded by JamA (**Figure 2**) but rational engineering failed to improve its interaction with JamB. We first constructed a library of PeACP variants by diversifying *peACP* (~250 bp) randomly using error-prone PCR such that each variant contains an average of two amino acid mutations. The mutant library of *peACP* was incorporated into a strain that co-expressed *jamA, jamB*, and *hspks1.* The strains expressing these mutants of PeACP were screened based on the titer of extracellular alkynes using a fluorogenic assay, which was previously established and employed for improving the activity of JamB.^15^ After screening ~900 clones followed by LC-HRMS confirmation, we identified six mutants (named as strains MS1_1 through MS1_6) with increased alkyne production efficiency (**Figure 6**). It is notable that although the best mutant strain only showed ~30% of the relative JamB activity compared to XZ1 (expressing *jamC)*, it had ~16-fold improved activity compared to the strain expressing the wild-type *peACP.*

**Figure 6.**
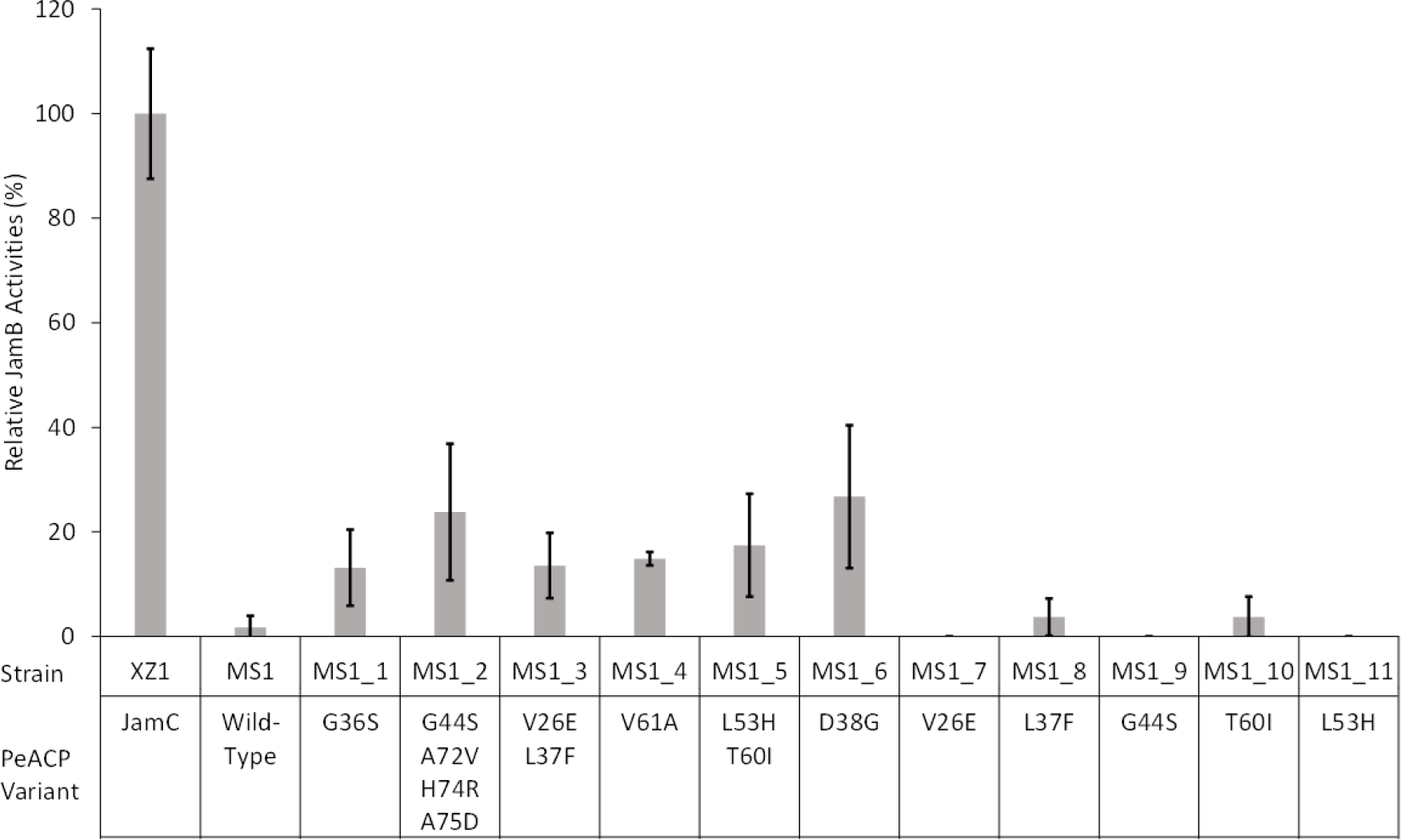
Relative JamB *in vivo* activities towards PeACP mutants. Error bars represent standard deviations from at least three experiments.

Sequencing analysis of these six *peACP* mutants showed that those from MS1_1, MS1_4, and MS1_6 contained one amino acid change, the mutants from MS1_3 and MS1_5 each had two mutations, and MS1_2’s mutant possessed four amino acid mutations (**Figure 6**). Single mutants corresponding to several amino acid changes found in MS1_3, MS1_5, and MS1_2 (except a few *C*-terminal residues) were constructed to evaluate the individual contributions of these mutations towards JamB recognition. None of these single mutants exhibited a significant positive effect (**Figure 6**), suggesting that some of these mutations have a synergistic effect on improving the relative JamB activity. It is worth noting that all five of the positive single mutations are either located right before PeACP helix 2 or at the beginning of PeACP helix 3, showing the hot spots on PeACP for protein engineering (**Figure. 7 and 8**). SDS-PAGE and biochemical analysis of each of these five PeACP mutants showed that none of these mutations significantly increased ACP expression or JamA activity (**Figures S1 and S5**), suggesting the improvements between the ACP and JamB recognition as the cause for the increase in compound **1** production. To better understand the molecular mechanism of these beneficial PeACP mutations, we mapped them to a modelled structure of the JamB-PeACP complex that was built using a previously reported method.^15^ These residues were predicted to locate at the interface of the two proteins, but the beneficial mutations could not be easily explained by electrostatic interactions (**Figure. 8 and S6**).

**Figure 7.**
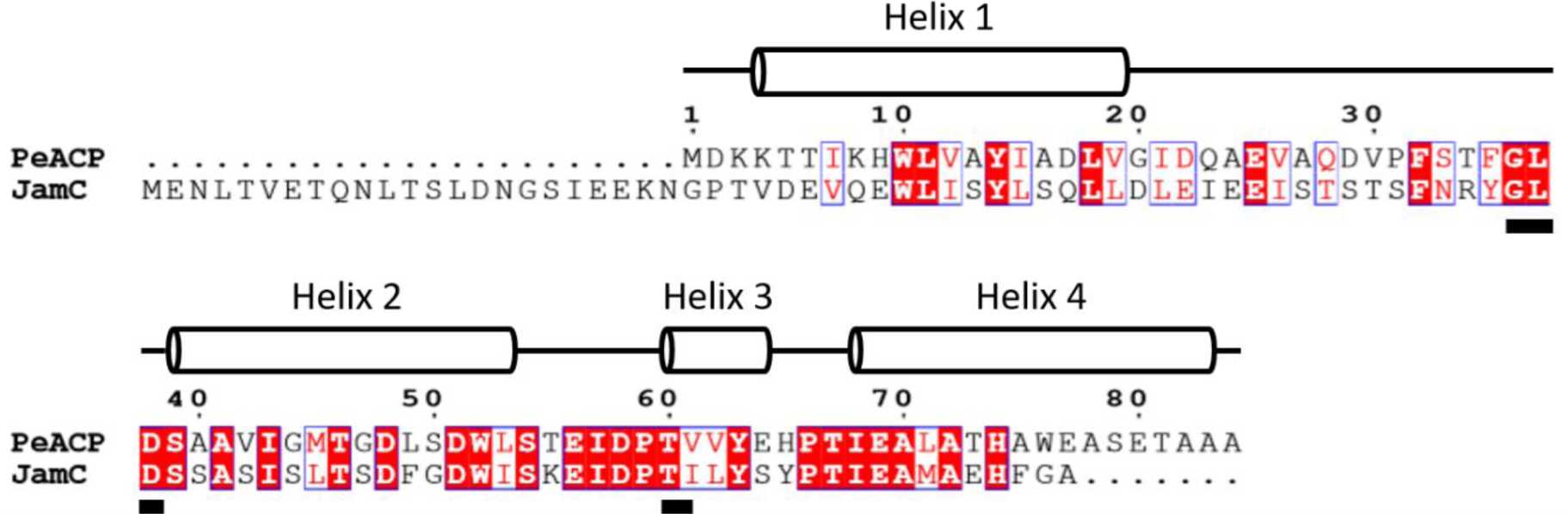
The amino acid sequence alignment of PeACP and JamC. The helical regions predicted from the modelled structures are indicated. The five PeACP residues as hot spots for providing JamB compatibility upon mutation are underlined.

**Figure 8.**
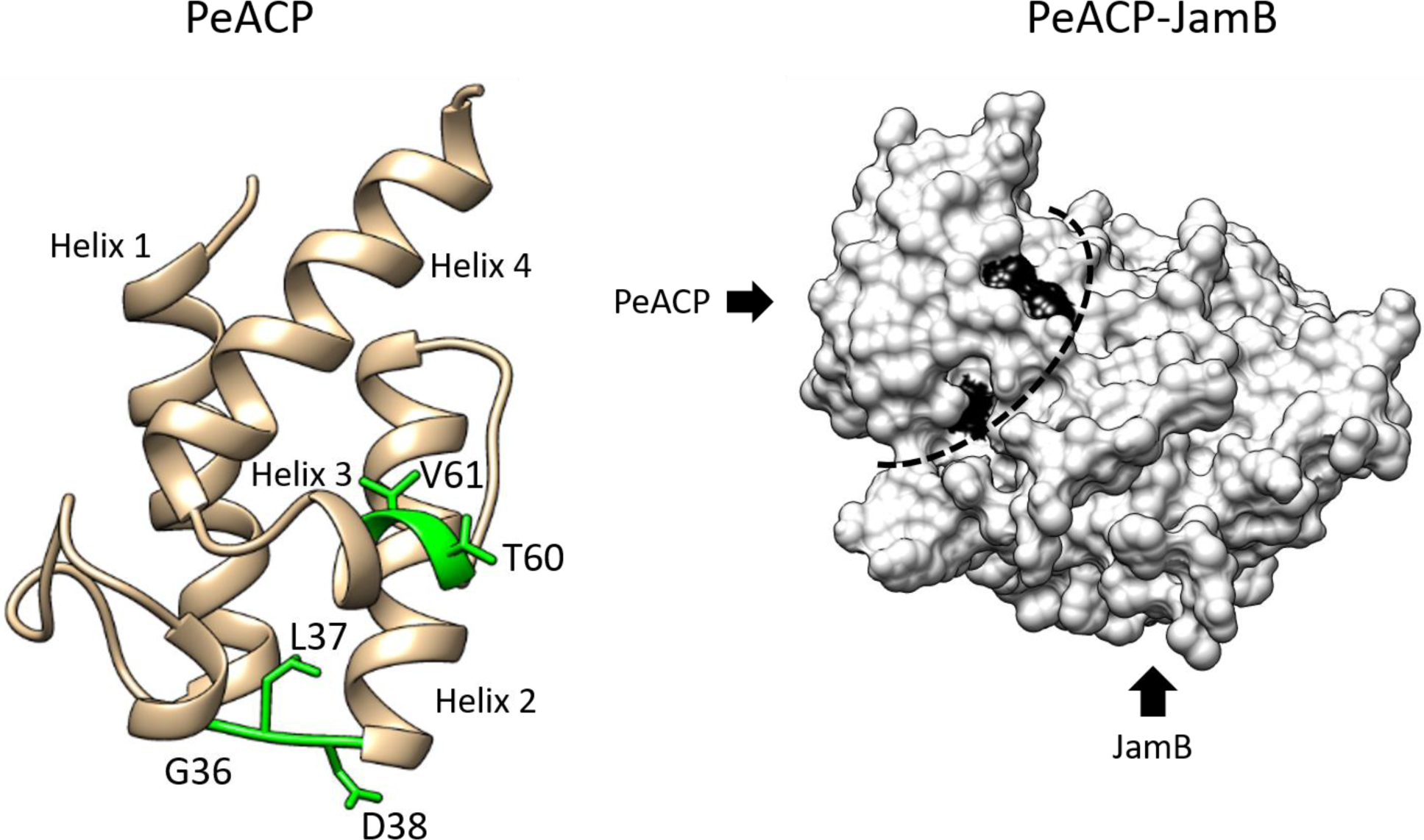
Modelled structures of PeACP and JamB. Left: A modelled ribbon diagram of PeACP. The five PeACP residues as hot spots for providing JamB compatibility upon mutation are shown as sticks. Right: Modelled surface structure of the JamB-PeACP complex. The two hot spots of the JamB-PeACP binding interface that were identified in the error-prone PCR and screening are marked.

## Conclusion

In summary, we have probed and improved the ACP-enzyme interactions within the alkyne biosynthetic machinery. Using both *in vitro* and *in vivo* analyses, we showed that the bi-functional desaturase/acetylenase (JamB) has a more stringent interaction with ACP than the acyl-ACP ligase (JamA). The recognition of JamB with non-cognate ACPs was further improved through ACP engineering. Specifically, rational engineering of TtuC based on the electrostatic interaction with JamB was explored, and a single charged amino acid mutation on helix 3 of TtuC improved the JamB recognition to a level comparable to the cognate ACP (JamC). Furthermore, we improved the interaction between JamB and PeACP through error-prone PCR and screening, and identified hot spots on PeACP that are important for interacting with JamB. This work has thus provided new insights into the critical role of ACP in the alkyne biosynthetic machinery, and showed the promise of protein engineering for generating a compatible heterologous ACP in alkyne biosynthesis to produce alkyne-tagged metabolites.

## Acknowledgements

We thank B. Pfeifer (University at Buffalo) for providing pBPJW144, and C. Bertozzi (Stanford) for providing the azido fluorogenic probe di-pegOF. This research was financially supported by the National Institutes of Health (Grant DP2AT009148), Alfred P. Sloan Foundation, and the Chan Zuckerberg Biohub Investigator Program.

